# Generation of lachrymatory factor synthase–suppressed onion (*Allium cepa* L.) by *Agrobacterium*-mediated gene transfer for CRISPR/Cas9 genome editing

**DOI:** 10.64898/2026.07.21.739049

**Authors:** Shoya Tamaru, Shinsuke Imai, Satoshi Watanabe, Tatsuya Ikegai, Sachie Kondo, Tomoko Igawa, Takahiro Kamoi

## Abstract

Lachrymatory factor, an irritating volatile with tear-inducing property, is produced when onion bulbs are cut or chopped. We aimed to generate onion plants with reduced lachrymatory factor synthase (LFS) activity via clustered regularly interspaced short palindromic repeats (CRISPR)/CRISPR associated protein 9 (CRISPR/Cas9) genome editing. Calli induced from primary roots were transformed with *Agrobacterium tumefaciens* carrying expression cassettes for CRISPR/Cas9, guide RNA, green fluorescent protein (GFP), and hygromycin resistance; callus lines that showed a high-frequency stable GFP expression were selected as “elite callus lines” that were suitable for transformation. Cleaved amplified polymorphic sequence (CAPS), heteroduplex mobility assay (HMA), and Sanger sequencing confirmed mutations introduced into the *LFS* gene, and plants were regenerated from the confirmed *LFS*-edited callus lines. The LFS enzyme activity in the leaves and bulbs of the *LFS*-edited plants was lower than that in control plants, while the *LFS*-edited plants exhibited severe growth abnormalities and failed to set seed, possibly due to long-term culture to maintain the elite callus line. The present study first demonstrated that onion genome editing, which modified a specific trait of onion, the reduction of LFS activity, was achieved. The results obtained opened the feasible way toward the final goal: the production of tear-free, higher health-functional onions.

## 1 Introduction

Onion (*Allium cepa L*.) is an important vegetable crop, ranking second in global production after tomato (FAOSTAT: Food and Agriculture Organization Corporate Statistical Database, 2024). Onion is notable for its lachrymatory effect that occurs when bulbs are cut. The lachrymatory factor, a volatile compound produced enzymatically from sulfenic acids, has been discovered to be synthesized by lachrymatory factor synthase (LFS) (Figure 1) (Imai et al., 2002). Low-lachrymatory onions with reduced LFS activity have been generated by RNA interference (RNAi) (Eady et al., 2008). The previous study showed that LFS suppression not only reduced the lachrymatory effect but also increased the amount of thiosulfinates, potentially contributing to higher health-functional properties and flavor. In the previous study, we established a model experimental system for evaluating onion bulb components using LFS-suppressed onions in comparison with normal onion (Eady et al., 2008). This system can estimate LFS from experimentally produced and quantified lachrymatory factor (LF) using the following procedures: sufficient alliinase is added to the substrate *E*-(+)-S-(1-propenyl)-L-cysteine sulfoxide (PRENCSO), and the generated intermediate 1-propenesulfenic acid is reacted with an onion reaction mixture containing LFS. Furthermore, the model system has identified the structures of thiosulfinates and sulfides that were elevated upon LFS suppression and revealed that some of these compounds possess inhibitory activity against enzymes such as cyclooxygenase-1 (COX-1), providing health-functional properties (Aoyagi et al., 2011; Aoyagi et al., 2021). To date, a production of another tear-free onion has been achieved through mutagenesis induced by a heavy-ion beam (Katoh et al., 2016). In the mutant lines, not only LF but also pyruvic acid were reduced due to repression of the alliinase-encoding gene expression, suggesting that thiosulfinate, which is involved in health-functional properties, was also reduced (Figure 1).

**Figure 1.**
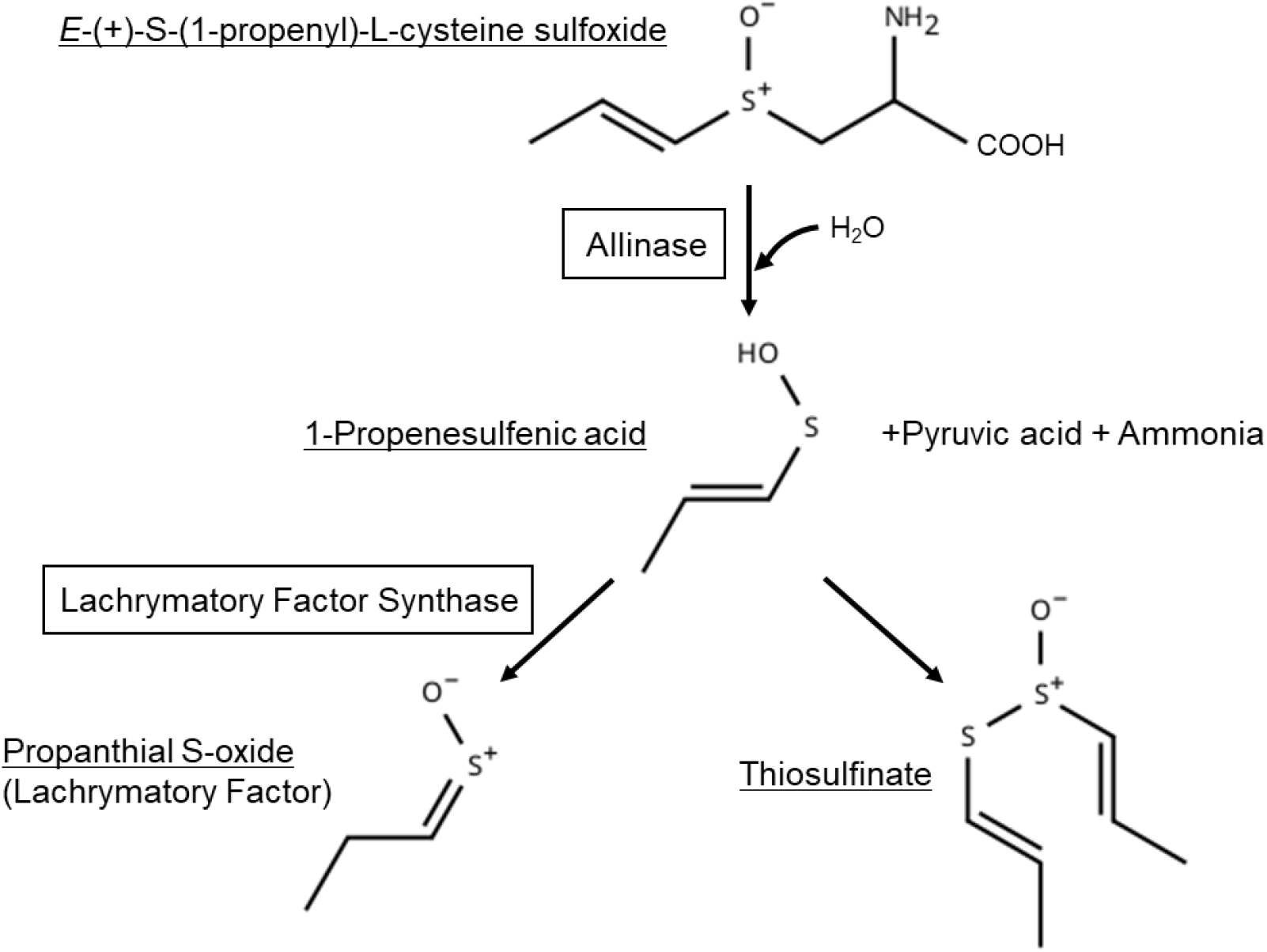
Biosynthetic pathway of lachrymatory compounds and thiosulfinates in onion. Alliinase catalyzes the substrate *E*-(+)-S-(1-propenyl)-L-cysteine sulfoxide (PRENCSO) to generate the intermediate 1-propenesulfenic acid, in addition to pyruvic acid and ammonia. Subsequently, lachrymatory factor synthase (LFS) catalyzes the conversion of this intermediate to produce lachrymatory factor (LF). At the same time, thiosulfinates, which are involved in health-functional properties, are also generated spontaneously.

As described above, our research group has conducted detailed, long-term analyses of onion-specific sulfur-containing compound metabolism and health-functional properties, with a particular focus on LFS. Building on these studies, LFS-specific suppression in onion is therefore expected to reduce the lachrymatory effect and enhance flavor and health-functional properties, improving suitability for raw consumption. To date, onions with low LFS activity have not yet been generated by conventional cross-breeding or occurred spontaneously.

Considering the biosafety and regulatory framework under the Cartagena Protocol, genome editing is a promising approach for precision breeding. In recent years, technologies such as clustered regularly interspaced short palindromic repeats (CRISPR)/CRISPR-associated protein 9 (CRISPR/Cas9) have been developed, enabling precise modification of target genes in plant genomes. However, onion has been recognized as one of the most recalcitrant crops for genetic transformation and genome editing due to its low transformation efficiency and further genotype-dependency. Consequently, only a few reports are available. Successful genome editing of the phytoene desaturase (*PDS*) gene in onion has been reported (Mainkar et al., 2023), whereas no reports targeting the genes that control onion-specific traits have been conducted to date. The present study reports for the first time the generation of onion in which the lachrymatory effect was reduced and flavor and health-functional components were increased through genome editing of the *LFS* gene.

## 2 Materials and Methods

### 2.1 Plant material

An onion cultivar ‘Senshu-Chu-Koudakaki’, which is an open-pollinated variety, was used in this study. In vitro seedlings were cultured under 12 h light at 25°C/12 h dark at 25°C to grow into larger than 15 cm or more. Bulb formation was then promoted by further culturing under 14 h light at 25°C/10 h dark at 25°C. Alternatively, large seedlings grown in vitro were transferred to soil, after which bulb enlargement was attempted following acclimatization. For acclimatization, plants transferred to soil were covered with plastic wrap to maintain humidity, and were gradually adapted to external environmental conditions over 1 week. After transfer to soil, plants were cultivated under 14 h light at 25°C/10 h dark at 25°C.

### 2.2 Construction of CRISPR/Cas9 vectors

Seven target sequences (#C2, #3, #9, #15, #R1, #R2, and #R3) for guide RNA (gRNA) were designed to target the regions proximal to the translation initiation or the enzymatically active sites within the *LFS* open reading frame (ORF). The positions of the targets and PCR primers are shown in Figure 2, and their nucleotide sequences are listed in Supplementary Tables 1 and 2.

**Figure 2.**
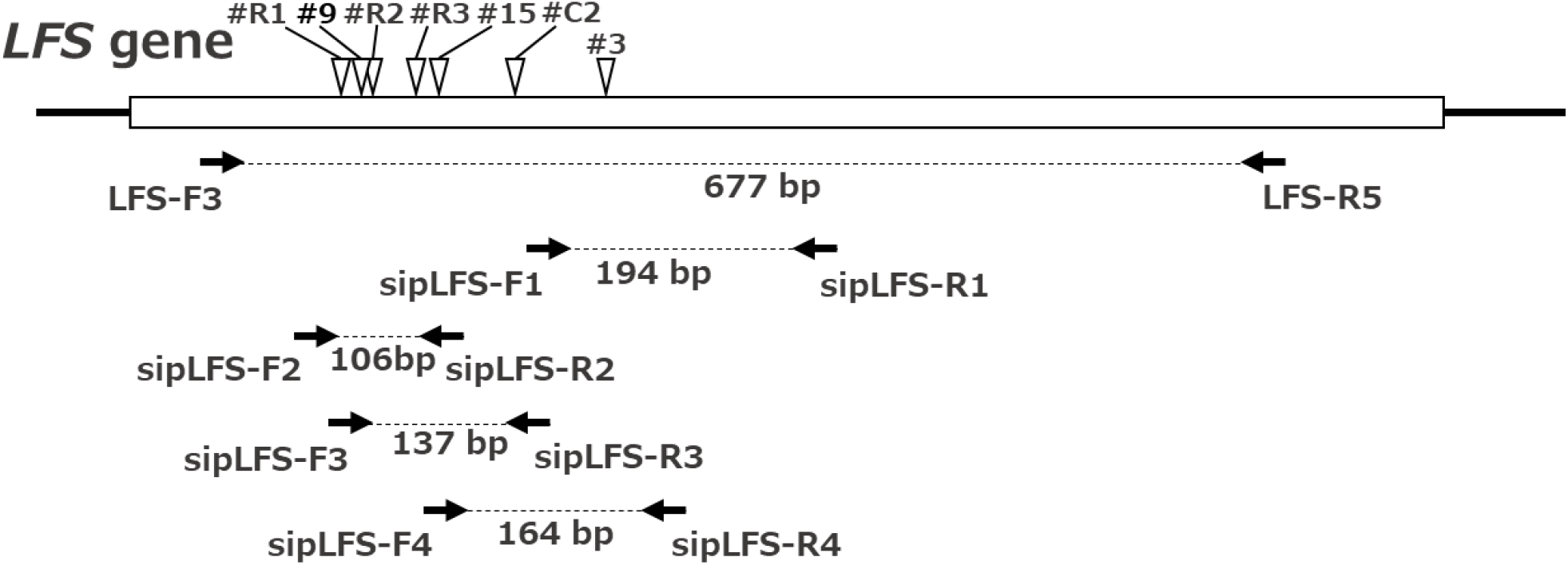
Positions of target sequences (triangles) and PCR primers (arrows) within the *LFS* gene (Accession no. AB089203.1). The amplicon size (bp) for each primer pair is indicated under the dashed line. Black vertical lines and a white box represent the untranslated regions and the intronless-ORF of *LFS*, respectively.

Two CRISPR/Cas9 vectors were constructed: pBI_Cas9-gRNA_HygR_GFP (carrying a single gRNA) (Figure 3A) and pBI_Cas9-gRNA_+3gRNA_HygR_GFP (carrying four gRNAs) (Figure 3B). The binary vector pBI-GW-NOS (Inplanta Innovations Inc., Yokohama, Japan) was used as the backbone. For both binary vectors, Cas9 and gRNA expression was driven by the mannopine synthase (MAS) promoter, which enables bidirectional transcription (Leung et al., 1991). To enhance translation of Cas9, the duplicated MAP-kinase activating protein 3 (dMac3) sequence (Aoki et al., 2014) was used. As selectable markers, hygromycin resistance gene (HygR) and enhanced green fluorescent protein (EGFP) were driven by the nopaline synthase (NOS) and cauliflower mosaic virus (CaMV) 35S promoters, respectively. The overall T-DNA structures of the two CRISPR/Cas9 vectors are shown in Figure 3.

**Figure 3.**
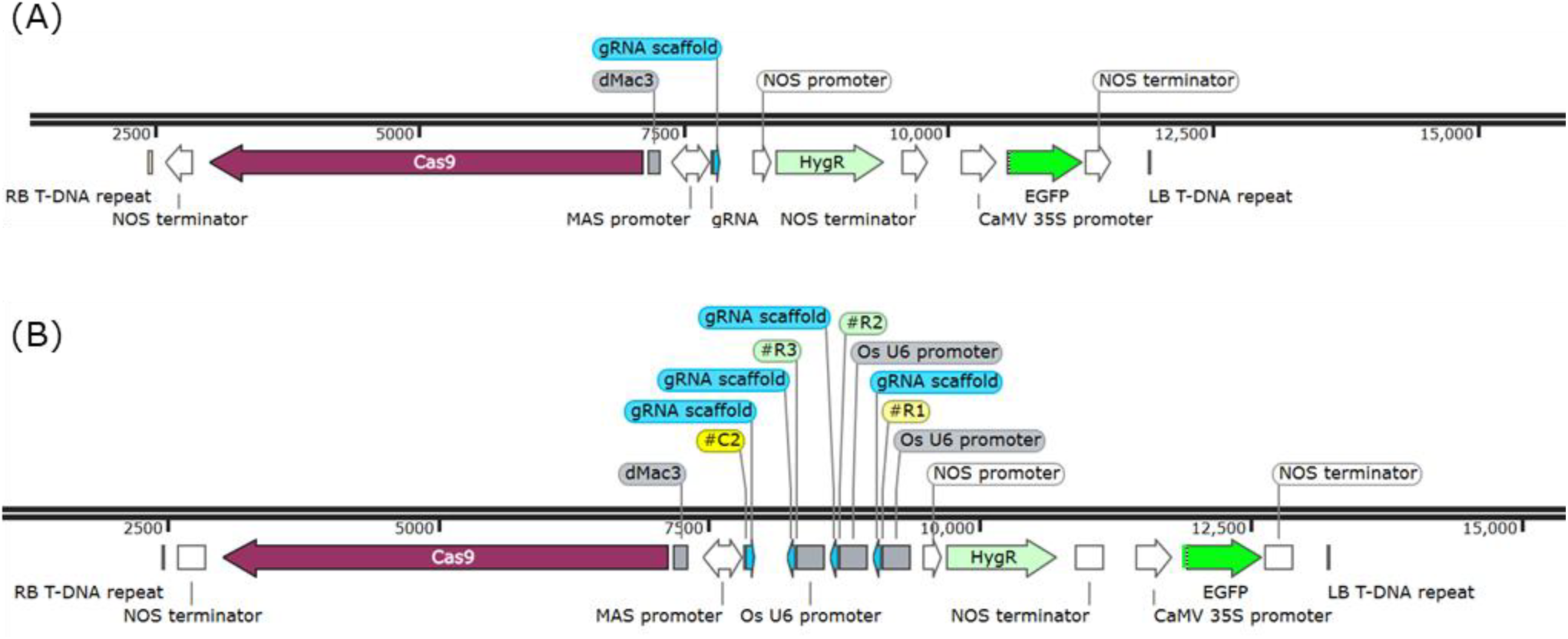
(A) T-DNA structure of CRISPR/Cas9 vector pBI_Cas9-gRNA_HygR_GFP, carrying a single gRNA (#C2, #3, #9, #15 each). (B) T-DNA structure of CRISPR/Cas9 vector pBI_Cas9-gRNA_+3gRNA_HygR_GFP, carrying four gRNA scaffolds (#C2, #R1, #R2, and #R3). LB, left border; RB, right border; NOS, nopaline synthase; dMac3, duplicated Mac3 translational enhancer; MAS, mannopine synthase; OsU6, Oryza sativa U6; HygR, hygromycin resistance gene; CaMV, cauliflower mosaic virus; EGFP, enhanced green fluorescent protein.

The constructed plasmids were introduced into *A. tumefaciens* strain LBA4404 as described by Ditta et al. (1980). *A. tumefaciens* culture and callus infection were performed as described previously (Kamata et al., 2011) with a minor modification; carbenicillin (250 mg•L^-1^) was used instead of cefotaxime (500 mg•L^-1^) in the selection and regeneration media. The compositions of the media used in the experiments are listed in Figure 4B.

**Figure 4.**
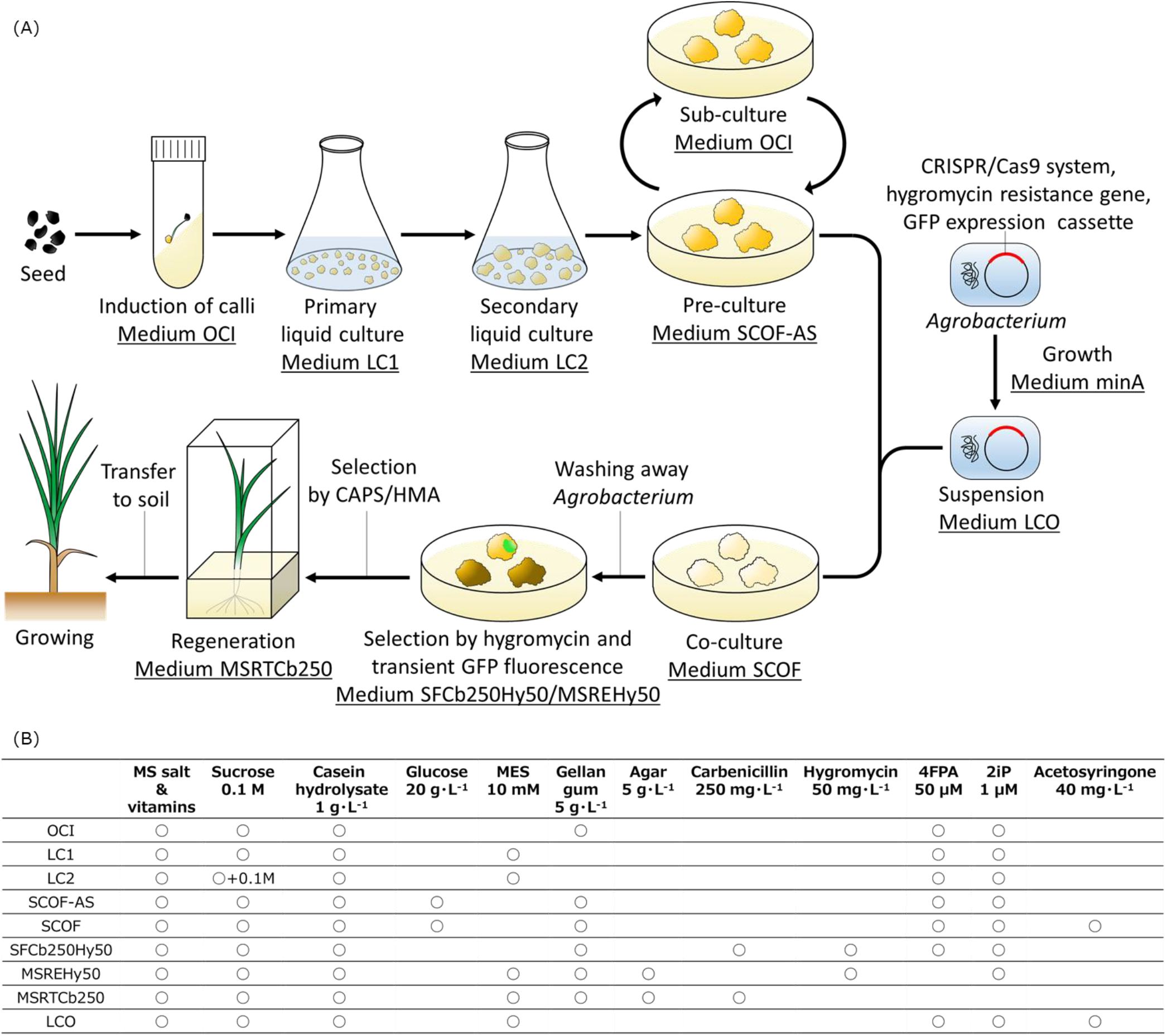
(A) Overview of the transformation and selection procedures to generate *LFS*-edited onion plants. GFP, green fluorescent protein; CAPS, cleaved amplified polymorphic sequence; HMA, heteroduplex mobility assay. (B) Composition of the culture media used in this study. MS, Murashige and Skoog medium; MES, 2-morpholinoethanesulphonic acid; 4FPA, 4-fluorophenoxyacetic acid; 2iP, N^6^(2-isopentenyl)adenine.

**Figure 5.**
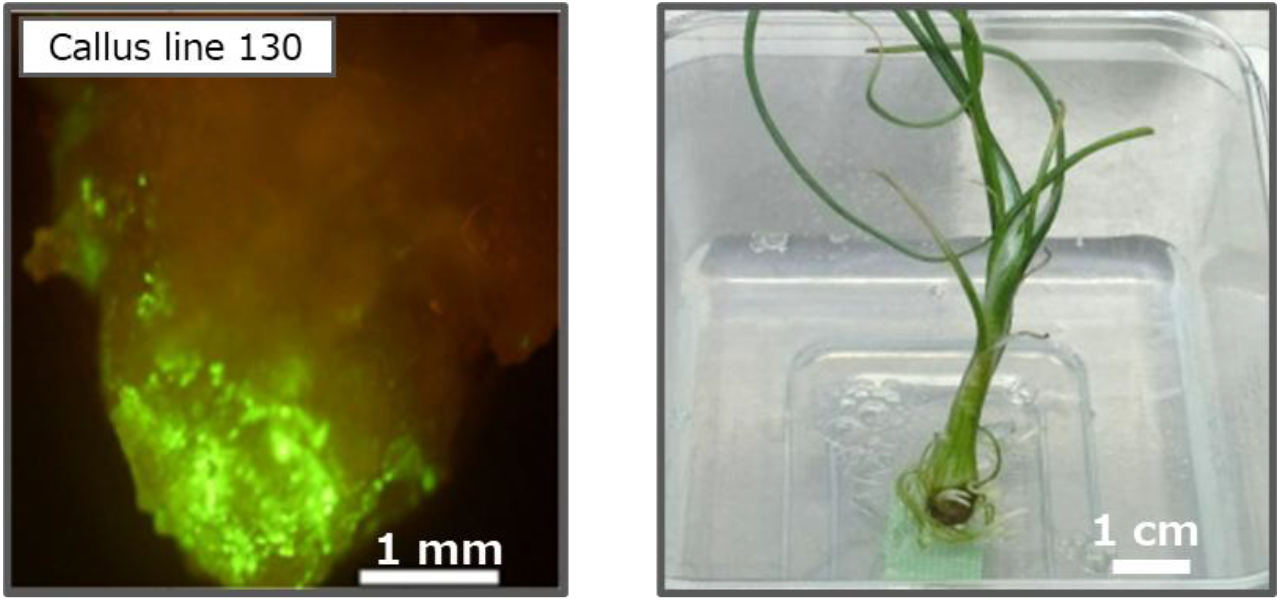
Stable GFP expression (left) and a regenerated plant (right) derived from callus line 130. Stable GFP expression was observed after one month from Agrobacterium infection.

### 2.3 Elite callus preparation

Callus induction and preparation for *Agrobacterium* infection were performed as described previously (Kamata et al., 2011). An overview of the procedure and the culture media used at each culture step is shown in Figure 4A. The cultured *Agrobacterium tumefaciens* cell suspension LBA4404 was prepared in minA solution (2.1 g •L^-1^ K_2_HPO_4_, 0.9 g •L^-1^ KH_2_PO_4_, 0.2 g •L^-1^ (NH_4_)_2_SO_4_, 0.1 g •L^-1^ trisodium citrate dihydrate, 0.2 g •L^-1^ MgSO_4_·7H_2_O, 2 g•L^-1^ glucose, and 400 mg •L^-1^ kanamycin). The compositions of all other media used in this study are listed in Figure 4B. For the selection of elite callus lines, Agrobacterium harboring a binary vector constructed using pBI-GW-NOS as the backbone vector, which carries a GFP expression cassette, was used for infection. The two vectors in Figure 3 were also used in the evaluations. After one month of culture, stable GFP expression was observed as described in section 2.4. The callus line exhibiting GFP expression at a higher rate with reproducibility was selected as the elite line.

### 2.4 Microscopy

GFP signal was observed using a Nikon SMZ18 fluorescence stereomicroscope with a filter cube (P2-EFL GFP-B or P2-EFL GFP-L) (NICON SOLUTIONS, CO., LTD., Tokyo, Japan). Calli with detectable GFP signal one month after transformation were used as transgenic plants for subsequent genetic analyses.

### 2.5 *LFS* mutation analysis of transgenic calli and plant lines

Genomic DNA (gDNA) was extracted from calli, leaves, and bulbs of regenerated plants using a DNeasy Plant Mini Kit (QIAGEN, Hilden, Germany). If a suitable recognition site by a restriction enzyme was present or absent in the target sequence, each CAPS or mobility assay (HMA) analysis was performed using primers listed in Supplementary Table 2. For both analyses, the nested PCR products amplified from the LFS-F3/LFS-R5 amplicon were used. Electrophoretic patterns were examined using the MultiNA microchip electrophoresis system (Shimadzu Corporation, Kyoto, Japan).

Mutation patterns were detected by Sanger sequencing or next-generation sequencing (NGS) of the partial LFS amplified with the primer set LFS-F3/LFS-R5 (Figure 2 and Supplementary Table 2) using KOD FX DNA polymerase (TOYOBO Co., Ltd., Osaka, Japan) at an annealing temperature of 55°C. Mutation frequencies were calculated as the proportion of reads carrying mutations per total number of reads (approximately 50,000–80,000 reads per plant).

### 2.6 LFS activity measurement

LFS activity was measured as described by Eady et al. (2008). To homogenize the samples, the harvested in vitro-grown leaves or bulbs were freeze-dried and ground into a powder. Each mixed suspension water containing the same weight of freeze-dried powder was centrifuged (20,400 ×g, 5 min, 4°C). The resulting supernatant was used as the LFS enzyme extract. Protein concentration was determined using a BCA Protein Assay Kit (Thermo Fisher Scientific, Waltham, MA, USA), according to the manufacturer’s instructions. Protein concentrations were adjusted uniformly among the samples, and relative LFS activities for each extract were measured.

## 3 Results

### 3.1 Selection of elite callus lines

Kamata et al. (2011) reported that calli with weak yellow autofluorescence under fluorescence microscopy with the GFP filter block showed higher Agrobacterium-mediated transformation efficiency. In this study, we further found that stable GFP expression rates differed among callus lines from individual seeds, even within the same open-pollinated variety (‘Senshu-Chu-Koudakaki’), showing strong genotypic dependence (Table 1). The stable GFP expression level observed in each callus line was reproducible across multiple experimental lots (Table 1). Although reducing totipotency during long-term maintenance of callus cultures is a well-known phenomenon, we confirmed that the regenerative abilities of these lines were maintained for more than one year through subculturing on OCI medium (Figures 4B and 5). Based on these results, selecting the elite callus line which exhibit higher GFP expression rate was important for subsequent Agrobacterium infection.

**Table 1.** Stable GFP expression rate in different callus lines.

| Callus line | Experiment No. |  |  |  |
| --- | --- | --- | --- | --- |
|  | 1 | 2 | 3 | 4 |
| 130 | 59.4% (329/554) | 20.5% (61/297) | 86.3% (113/131) | 66.0% (157/238) |
| 132 | 12.9% (60/465) | 9.8% (49/499) | - - | - - |
| 134 | 20.0% (55/275) | 28.7% (80/279) | - - | - - |
| 136 | 17.9% (122/683) | 41.3% (71/172) | - - | - - |
| 145 | 7.1% (39/553) | 18.4% (73/397) | 4.8% (6/126) | 4.2% (12/285) |
| 147 | 8.2% (29/353) | 10.6% (91/857) | 2.0% (3/150) | 3.4% (11/326) |

### 3.2 *LFS* mutation analysis of the transgenic calli and regenerated plants

The results of genome editing by five different gRNAs are summarized in Table 2. Among 483 transgenic callus lines stably expressing GFP, *LFS* mutations were detected by CAPS and HMA analyses in 101 lines in total. Consequently, the mutations were detected in callus lines targeted by #C2, #15, or #3-harboring single gRNAs and #R1, R2, R3, and C2-quadruple gRNAs, but not by #9-harboring single gRNA.

**Table 2.** CRISPR/Cas9-induced mutations in callus lines and regenerated plants by gRNA.

| Target | No. of callus |  | No. of regenerated plants |  |
| --- | --- | --- | --- | --- |
|  | Analysed | Edited | Analysed | Edited |
| #R1, R2, R3, C2 | 55 | 22 | 15 | 4 |
| #C2 | 120 | 28 | 7 | 7 |
| #15 | 258 | 47 | 8 | 8 |
| #9 | 33 | 0 | 0 | 0 |
| #3 | 17 | 4 | 0 | 0 |

In total, 71 regenerated plants were obtained from the mutation-positive callus lines, while no plants were obtained from the calli with mutations in the #3 target sequence. Of these regenerated plants, 30 plant lines exhibiting fine growth were subjected to NGS analysis to characterize the mutation patterns and the frequencies (Table 3). Consequently, *LFS* mutations were confirmed in 19 regenerated plant lines. Among these, five plant lines (P2, P20, P40, 130, and 464) with mutation frequencies exceeding 50% were identified, and 3-7 mutation patterns were detected in each line (Table 3 and Figures 6, 7A).

**Table 3.** The number of *LFS*-mutated plants and their mutation frequencies.

| Target | No. of *LFS*-edited plants |  |  |  |  |  |  |  |  |  |
| --- | --- | --- | --- | --- | --- | --- | --- | --- | --- | --- |
|  | Mutation frequencies by NGS (%) |  |  |  |  |  |  |  |  |  |
|  | 0-10 | 10-20 | 20-30 | 30-40 | 40-50 | 50-60 | 60-70 | 70-80 | 80-90 | 90-100 |
| #R1, R2, R3, C2 | 11 | - | - | 1 | - | - | **1** | - | **1** | **1** |
| #C2 | - | - | 3 | 2 | 1 | **1** | - | - | - | - |
| #15 | - | 4 | 2 | 1 | - | **1** | - | - | - | - |

Target sequences included in the gRNA were represented in the left columns. The number of plants showing each mutation frequency range determined by NGS is indicated in the table. The number of plant lines with a mutation frequency of more than 50% is labeled in red.

**Figure 6.**
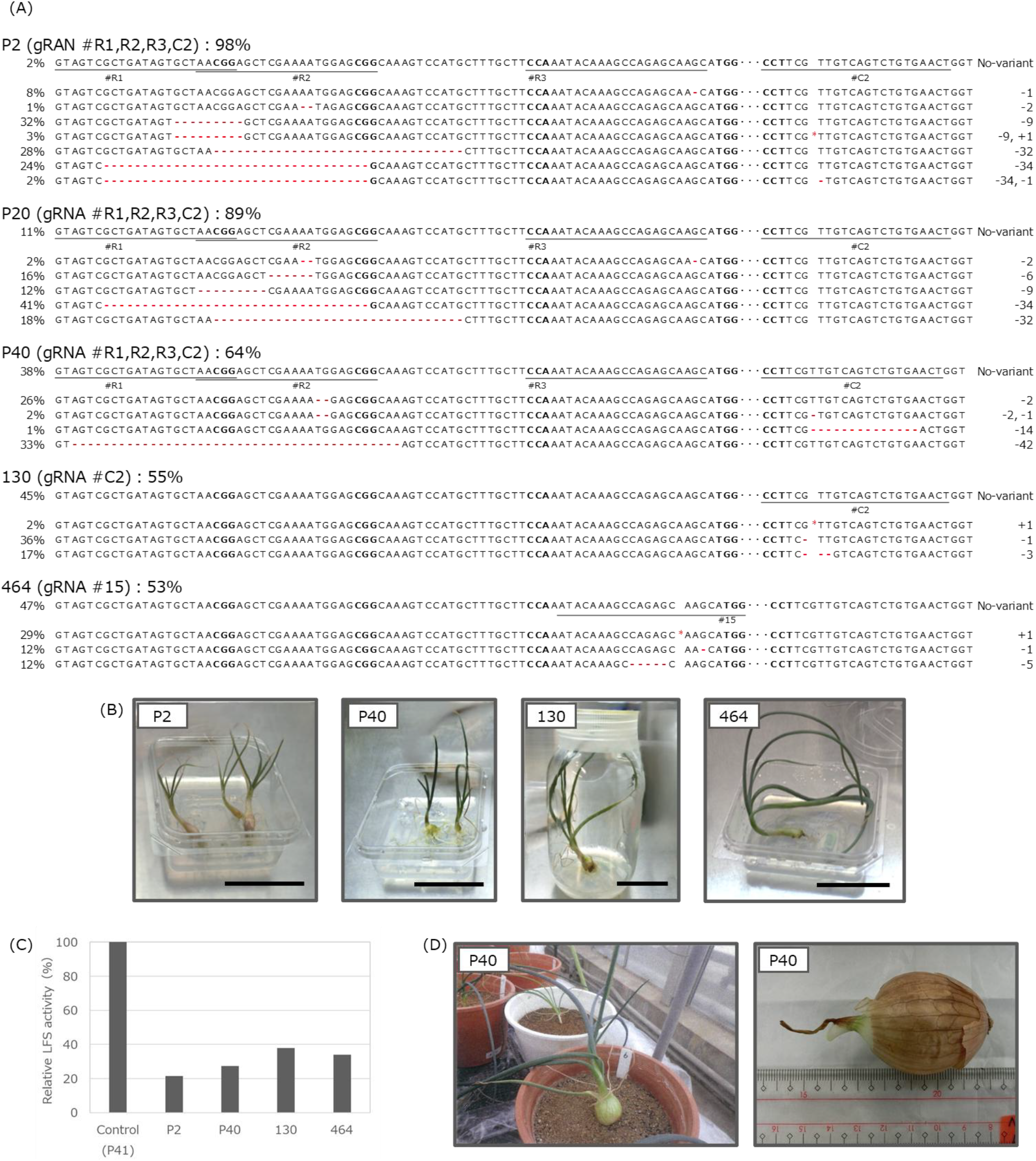
The *LFS* mutant *(lfs*) plant lines and their LFS enzyme activities. (A) Mutation patterns and the frequencies in each *lfs* plant line (T0). Target sequences are underlined respectively. Bold nucleotides in each target indicate the PAM sequence. Red hyphen (-) and asterisk (*) denote the nucleotide deletion and insertion, respectively. The numbers on the right indicate the number of inserted (+) and/or deleted (-) nucleotides detected in each sequence. (B) In vitro grown-regenerated plant lines P2, P40, 130, and 464 (from left to right). Bar = 5 cm. (C) Relative LFS enzyme activity in leaves of the *lfs* plant lines. The average activity (%) from two measurements in each sample is indicated. (D) Growth of plant line P40 during soil cultivation (left) and the harvested bulb (right).

**Figure 7.**
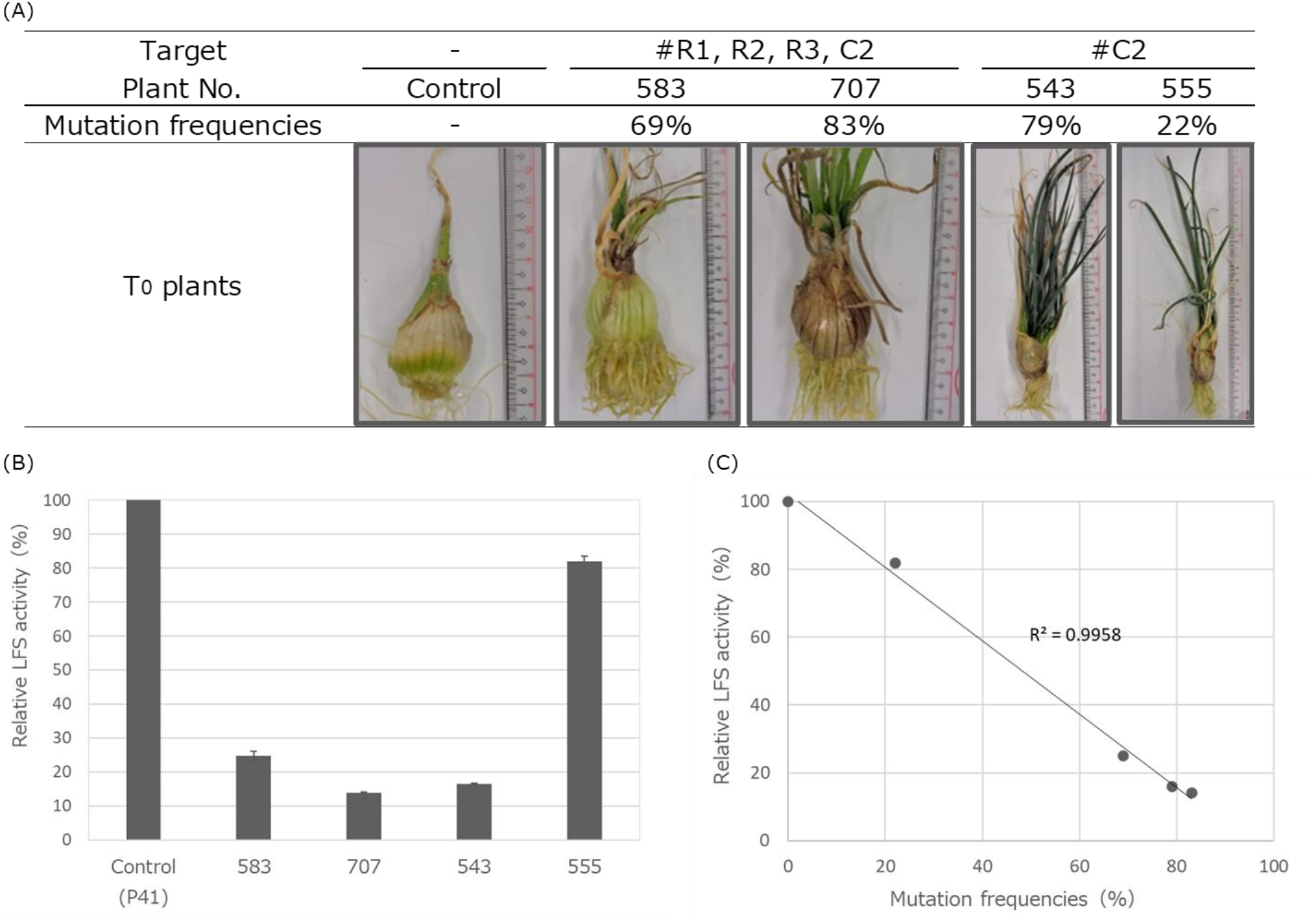
In vitro bulb formation for evaluation of LFS activity. (A) In vitro-formed bulbs of the *lfs* plants and their morphologies. Targets for obtaining the genome-edited plants are indicated at the top of each photo. The mutation frequencies determined by NGS are indicated under the T0 plant numbers. (B) Relative LFS activity in bulbs of T0 *lfs* plant lines. The average relative activity (%) from three measurements in each sample is indicated. The vertical bars on each graph represent the standard deviation. (C) Relationship between mutation frequency and relative LFS activity in bulbs from T0 *lfs* plants. The point at 0% mutation frequency/100% LFS activity is derived from the non-transgenic control.

### 3.3 LFS enzyme activity in the leaves of *lfs* plants

LFS activities in the leaves of the four mutant plant lines (P2, P40, 130, and 464) were measured (Figure 6C). Plant P41 was a line derived from a transgenic callus line introduced with gRNAs for #R1, R2, R3, and C2. Because NGS confirmed no *LFS* mutations, it was used as the experimental control. As a result, relative LFS activities in all four *lfs* plant lines were reduced to less than 40% of the control plant P41. The most reduced activity was detected in the P2 plant line, in which the highest mutation frequency was detected (Table 3 and Figure 6A). On the other hand, no clear correlation between the mutation frequencies and the reduction rate was observed in LFS activity from the leaves (data not shown).

These four in vitro grown *lfs* plants were transplanted to soil to obtain sufficient size of bulbs and progeny seeds. However, all the plants exhibited considerably retarded growth than control seedlings, and plants P2, 130, and 464 eventually died. In the case of plant P40, it toppled during cultivation, so we needed to harvest the bulb with approximately 20 mm in diameter, which was too small (Figure 6D). Consequently, the bulbs subjected to vernalization died during cold storage due to rot, and no progeny seeds were obtained.

### 3.4 LFS enzyme activity in the freeze-dried bulbs derived from *lfs* plants

As soil cultivation of P2, P40, 130, and 464 plants was unsuccessful, we attempted in vitro bulb formation instead. We newly obtained the T0 *lfs* plant lines 543, 555, 583, and 707 (Figure 7A) by the procedures described above. These in vitro plants were cultured in large vessels containing MSREHy50 medium (Figure 4B) under long-day conditions (18 h light at 24°C/6 h dark at 20°C).

All the newly obtained *lfs* plants succeeded in bulb formation, and especially plant 707 produced a bulb with approximately 30 mm in diameter and about 10 g in weight (Figure 7A), although the bulb sizes were not comparable to those produced in general cultivation of wild-type onions. In this experiment, the control plants, regenerated from non-transgenic calli, were also cultured under identical conditions, and these also exhibited small bulb formation as well. In contrast, seedlings obtained directly from seeds, without a callus phase, grew and developed bulbs exceeding 100 mm in diameter under the same in vitro conditions.

Next, the LFS activities in the obtained bulbs were measured. All samples showed lower LFS activities than the non-transgenic control (Figure 7B). Unlike the result of LFS activities in the leaves (Figure 6C), LFS activity in the bulbs showed a correlation between LFS reduction rate and mutation frequency (Figure 7C).

## 4 Discussion

### 4.1 Callus line suitable for Agrobacterium transformation

In *Agrobacterium*-mediated transformation experiments using onion callus, high efficiencies of both plant regeneration and transformation are important. Tanikawa et al. (1998) reported that plant regeneration efficiency from callus differs among onion cultivars, and Kamata et al. (2011) reported that calli with weak autofluorescence show higher transformation efficiency than those with strong autofluorescence, even when derived from the same cultivar. Based on these findings, we used ‘Senshu-Chu-Koudakaki’ as an appropriate cultivar in the present study, and selected calli showing weak autofluorescence for transformation. We further confirmed strong genotype dependency for transformation efficiency by comparing calli derived from individual seeds. Any morphological differences between calli with low and high transformation efficiencies were not distinguished at the stereomicroscopy level. Interestingly, elite callus lines with high GFP expression rates maintained the trait even after a long culture period with repeated subculturing. Therefore, preliminary evaluation with a visible marker is a critical step for onion transformation mediated by Agrobacterium. Future analyses, such as RNA-seq, proteome, and phytohormone measurement, would accumulate molecular and physiological insights into differences in recalcitrant/amenable cell lines, contributing to the easy selection of elite lines.

### 4.2 Selection of optimal target sequence for Cas9-inducible indel mutation

In this study, *lfs* mutant plants were successfully obtained by *LFS-*targeting genome editing. We designed seven target sequences for gRNAs considering positions that influence translation or enzyme activity. Consequently, mutations were detected with #3, #15, #C2, #R1, and #R2-gRNAs, whereas no mutations were detected with #9 or #R3-gRNAs. The difference in mutation frequencies (Figure 6A) may depend on the Cas9-inducible mutation efficiency on the sequence. In recent years, AI-based gRNA design programs such as DeepSpCas9 (https://deepcrispr.info/) (Kim et al., 2019) have been developed. The DeepSpCas9 score of each target sequence used in this study showed that targets with scores over 40 caused mutations, whereas two of the three targets with less than 40 produced no detectable mutations, except for target #3 (Supplementary Table 3). Especially, #C2 with the highest score provided a *lfs* mutant with more than 50% mutation frequency, and it seemed that the combination with #R1 and #R2 enhanced the mutation frequency (Table 3). As the outcomes in this study showed a rough correlation between the DeepSpCas9 score and mutation efficiency, the employment of recent prediction tools in target sequence settings is effective.

### 4.3 Achievement of LFS activity reduction in the *lfs* plants

Based on our past discovery of the LFS enzyme for the production of lachrymatory factor, the malfunction of the LFS is an ideal scheme for the generation of tear-free onions with higher health-functional properties (Imai et al., 2002; Eady et al., 2008). Additionally, precision breeding is highly demanded. In this study, the *lfs* plant lines with over 50% mutation frequencies exhibited reduced LFS activities, indicating that the designed scheme was achieved as expected. But unexpectedly, the *lfs* plants did not grow after acclimatization; therefore, the normal size of bulbs and the progeny seeds were not obtained.

The use of mature seeds for callus induction has an advantage in year-round material and easier manipulation compared to the previous study using calli induced from immature embryos (Eady et al., 2000). In addition, all elite callus lines subjected to long-term subculture for 3 years or longer were able to regenerate the plants. However, very poor growth in soil after acclimatization resulted in severely stunted bulb development, with bulbs not exceeding approximately 4 cm in diameter. Non-transgenic plants derived from non-elite callus lines maintained for 3 years longer also produced bulbs not exceeding approximately 4 cm in diameter (data not shown). Moreover, the defects in bulb formation with morphological abnormalities in roots and leaves in soil cultivation have been observed in all plants derived from different elite callus lines, which were induced from different seed genotypes. Even in the in vitro-grown plants that succeeded in larger bulb formation (Figure 7A), abnormalities such as bushy leaves and non-elongating roots were also observed. Given these results, the observed poor growth was unlikely to be attributable to off-target effects of the gRNAs, in addition to the particular seed genotype used here, and outcomes from LFS reduction. It was considered that somaclonal variation accompanied by epigenetic modifications could have occurred (Kaeppler et al. 2000; Debnath and Ghosh, 2022), although the causes of poor growth under soil cultivation remain unclear. In onion, to date, long-term callus culture is an essential step that has been used for *Agrobacterium*-mediated transformation (Eady et al., 2000; Tanikawa et al., 2006; Kamata et al., 2011). To circumvent the unexpected plant growth retardance and vulnerability observed in this study, the development of new genome editing approaches that do not require a callus phase is highly desirable. Recently, *in planta* transformation in which the shoot apical meristem is targeted for CRISPR/Cas9 gene transfer has succeeded in the generation of genome-edited plants (Imai et al., 2020; Kumagai et al., 2022; Sasaki et al., 2025; Sebiani-Calvo et al., 2024). With our scheme for the production of tear-free onions, the employment of in planta transformation technologies would be the subsequent potential strategy.

## 5 Conclusions

In this study, we succeeded in generating the *LFS* genome-edited onion plants for the first time. This was achieved by *Agrobacterium*-mediated *CRISPR*/*Cas9* and gRNA transfer through selecting elite callus lines with high transformation efficiency. For genome editing, we used the *Agrobacterium*-mediated gene delivery described previously (Kamata et al., 2011), in which calli induced from primary roots serve as explants. As expected, the LFS enzyme activity was reduced in the bulbs from genome-edited plants, thereby providing a proof-of-concept for the generation of tear-free onions.

However, the problem in the growth of regenerated plants and the bulb formation in soil cultivation, likely associated with long-term callus culture, remains. The alternative genome editing methods for onion that do not require a callus phase would enable the desirable tear-free onion production with normal growth.

## 6 Acknowledgments

We are grateful to Drs. Toki, Hirose, Endo, and Saika for providing the CRISPR/Cas9 construct and the gRNA scaffold, and to Prof. Shimada for providing the dMac3 construct. We also thank Prof.

Ezura for valuable advice on genome editing, Drs. C. Eady, F. Kenel, and S. Davis for advice on onion transformation, and Dr Makabe for technical assistance in vector constructions.

## 8 Author contributions

ST: Conceptualization, Methodology, Investigation, Formal analysis, Writing – original draft. TK: Conceptualization, Methodology, Investigation, Formal analysis, Writing – review and editing, Supervision, Project administration, Funding acquisition. SI: Conceptualization, Methodology, Investigation, Formal analysis, Writing – review and editing, Supervision, Project administration. TI: Conceptualization, Methodology, Investigation, Formal analysis. SW: Formal analysis. SK: Formal analysis. TI: Conceptualization, Methodology, Investigation, Formal analysis, Writing – review and editing, Supervision, Project administration, Funding acquisition. All authors contributed to the article and approved the submitted version.

## 10 Data availability statement

The original contributions presented in the study are included in the article and its Supplementary Material. Further inquiries can be directed to the corresponding author.

