## Supplementary Material for "Generation of lachrymatory factor synthase–suppressed onion (*Allium cepa* L.) by *Agrobacterium*-mediated gene transfer for CRISPR/Cas9 genome editing"

### 1 Supplementary Material

#### 1.1 Supplementary Tables

Supplementary Table 1. Target sequences for *LFS* genome editing.

| Target | Sequence (5′ → 3′) PAM |
|---|---|
| #R1 | GTAGTCGCTGATAGTGCTAA**CGG** |
| #R2 | AACGGAGCTCGAAAATGGAG**CGG** |
| #R3 | **CCA**AATACAAAGCCAGAGCAAGC |
| #C2 | **CCT**TCGTTGTCAGTCTGTGAACT |
| #3 | GGTTGTGTTCGCTACGTTAA**AGG** |
| #9 | GTGCTAACGGAGCTCGAAAA**TGG** |
| #15 | ATACAAAGCCAGAGCAAGCA**TGG** |

PAM (protospacer adjacent motif) sequences are indicated in red letters.

Supplementary Table 2. Primers used in this study.

| Primer | Sequence (5′ → 3′) | Analyzed target | Analysis |
|---|---|---|---|
| LFS-F3 | CAATTCAGACTCACATTACGTTAT | #R1, R2, R3 | NGS |
| LFS-R5 | ACACACAACACTCAGTCTTAC | | |
| sipLFS-F4 | CAAATACAAAGCCAGAGCAAGC | #C2 | Nested PCR for HMA |
| sipLFS-R4 | AAATTCCTCTTCTATTGGGTGCAT | | |
| sipLFS-F1 | GCCTTCGTTGTCAGTCTGTGAACT | #3 | Nested PCR for CAPS |
| sipLFS-R1 | CCGTGTAATCCTCGTACCCTGTAA | | |
| sipLFS-F2 | AAATCCTGGTGCACCTGC | #9 | Nested PCR for CAPS |
| sipLFS-R2 | CCATGCTTGCTCTGGCTTTG | | |
| sipLFS-F3 | TAGTCGCTGATAGTGCTAACGG | #15 | Nested PCR for CAPS |
| sipLFS-R3 | TGACAACGAAGGCATGACC | | |

Supplementary Table 3. DeepSpCas9 scores of targets used in this study

| Target | DeepSpCas9 score | Mutation detected |
|---|---|---|
| #C2 | 81.408 | ○ |
| #R2 | 56.103 | ○ |
| #15 | 50.624 | ○ |
| #R1 | 42.270 | ○ |
| #9 | 31.451 | × |
| #3 | 15.793 | ○ |
| #R3 | 12.949 | × |

Detectable mutation outcome for each target is indicated as ○(detected) or × (not detected).
